# JAK2 is highly expressed by cyst-lining cells and regulates cystogenesis

**DOI:** 10.1101/378919

**Authors:** Foteini Patera, Alex Cudzich-Mardy, Zhi Huang, Maria Fragiadaki

**Author notes:** **Correspondence to:** Dr Maria Fragiadaki, Academic Nephrology Unit, Department of Infection, Immunity and Cardiovascular Disease, University of Sheffield, Beech Hill Road, S10 2RX. These authors contributed equally (Foteini Patera, Alex Cudzich-Mardy, Zhi Huang).

## Abstract

Dysregulated JAK/STAT signalling is implicated in polycystic kidney disease, which is a common genetic disease leading to renal failure. However, the mechanisms underlying JAK/STAT-mediated cystogenesis arepoorlyunderstood.TheroleofJAK2wasinvestigated immunohistochemically in a murine model of cystic disease (Pkd1^nl/nl^). In the normal kidney, JAK2 is restricted to tubular epithelial and vascular cells with lesser staining in bowman’s capsule and is undetectable in the interstitium. By contrast, in the diseased kidney JAK2 appears stronger in cyst-lining cells when compared to normal tubules and appears mislocalised in the interstitium. Given that JAK2 is a major tyrosine kinase activating JAK/STAT, we considered whether its inhibition can attenuate cystic growth *in vitro*. To assess this we used curcumin, a natural phytochemical, which significantly reduced JAK2 levels and STAT3 activity. Consistently, reduced JAK2/STAT3 activity was correlated with reduced cystic growth of cystic cells in three-dimensional cyst assays. Taken together, our results suggest that JAK2 is a key signalling molecule that functions to inhibit cystic growth in cystic tubular epithelial cells, thus providing the foundation for its development as a novel therapeutic in polycystic kidney disease.

## INTRODUCTION

Autosomal dominant polycystic kidney disease (ADPKD) is a devastating multi-organ disease, affecting over 9 million people worldwide, lacking a cure. It accounts for approximately 5% of patients with renal failure. Genetically, it arises predominately due to mutations in the Pkd1 or Pkd2 genes, encoding for the polycystin-1 and polycystin-2 proteins. ADPKD is characterised by the progressive growth and enlargement of renal cysts, typically leading to renal failure by middle age. Mutations in polycystins also cause a strong vascular phenotype associated with hypertension and aneurysms ^1,2^ which together with cysts in other organs, shows that ADPKD is a systemic disease. The complexity of changes in ADPKD have made the identification of a cure difficult. Currently Tolvaptan, a V2-vasopressin receptor antagonist, is the only approved therapy; however, it carries significant side effects precluding its use in the majority of patients ^3^. Therefore, a major biomedical challenge is to identify central druggable pathways for the treatment of ADPKD.

JAK-STAT is an evolutionarily conserved pathway misregulated in a number of conditions such as cancer, arthritis and other inflammatory diseases ^4^. We, and others, have previously shown that this pathway is abnormally activated in, and contributes to, ADPKD ^5-9^. The pathway becomes activated by a series of tyrosine phosphorylation steps mediated by one of the four kinases JAK1-JAK3 or Tyk2. Because of their involvement in multiple diseases, several JAK inhibitors have been developed over the years, including the approved Ruxolitinib, Tofacitinib and Baracitinib ^10-13^. Moreover, additional JAK inhibitors are in clinical trials, making JAK-STAT a therapeutically tractable pathway. Combining our previous and current work with that of other groups we predict that JAK-STAT inhibition may be of therapeutic benefit in ADPKD.

Given that ADPKD is a complex multiorgan disease we reasoned that selective inhibitors might not be beneficial in the initial stage of disease when a plethora of responses are raging. Without excluding the possibility that later, when the disease is more controlled, perhaps a more selective JAK inhibitor might be an effective and safe strategy for maintenance therapy.

Curcumin (diferuloylmethane) is a phytochemical ^14^ that exhibits actions against a number of pathways including the JAK-STAT thus exhibiting strong antioxidant, anti-tumour and anti-inflammatory pharmacological properties ^15^. Importantly, no appreciable side-effects have been reported making it an attractive compound to inhibit JAK-STAT. Because its low toxicity but potent activity, curcumin has been tested in ADPKD murine models of disease where it was able to limit the growth of cysts ^16^ and is currently being trialled in young patients and children with ADPKD (clinicaltrials.gov). We therefore used curcumin to inhibit JAK-STAT activity in ADPKD-derived renal cells. In doing so we uncovered that curcumin inhibits JAK-STAT and reduces the rate of growth of cysts by inactivating JAK2 in renal tubular epithelial cells. Collectively our data suggest a central role of JAK2 in the development and growth of cysts in polycystic kidney disease.

## RESULTS

### JAK2 is highly expressed by renal cystic epithelial cells *in vivo*

To investigate whether JAK2 may be involved in the development of cysts *in vivo*, we stained kidney sections from the Pkd1^nl/nl^ mouse model ^17^ and associated wild-type littermate control kidneys with anti-JAK2 antibodies. Kidneys from 5 weeks old mice, with intermediate disease, and 10 weeks old with advance disease were studied, as previously described ^5^. We found that JAK2 is widely expressed by cortical renal epithelial cells and its expression appears predominately cytoplasmic with no detectable expression by interstitial fibroblasts or in the interstitium (Fig. 1, top). During early stages of ADPKD, JAK2 is strongly expressed by renal cystic tubules, and to a lesser extent by non-cystic, otherwise normal, tubular epithelial cells (Fig. 1, middle). As the disease advances at 10 weeks of age (as evidenced by increased number of larger cysts and diffuse interstitial expansion), JAK2 is strongly expressed in cystic epithelial cells and this expression appears to be stronger in cystic tubules than that observed in healthy non-cystic tubules. Some expression is detected in interstitial cells (Fig 1, bottom) which is absent in healthy kidneys (Fig. 1 top), suggesting JAK2 expression in the interstitium is mislocalised. JAK2 is expressed in vessels with a lesser expresion by bowman’s capsule cells (Figure 1, bottom). Expression of JAK2 in vessels remains strong as disease progresses (Fig. 1, bottom). Hence, JAK2 expression is temporally and spatially coincident with cyst development in the ADPKD kidney.

**Figure 1:**
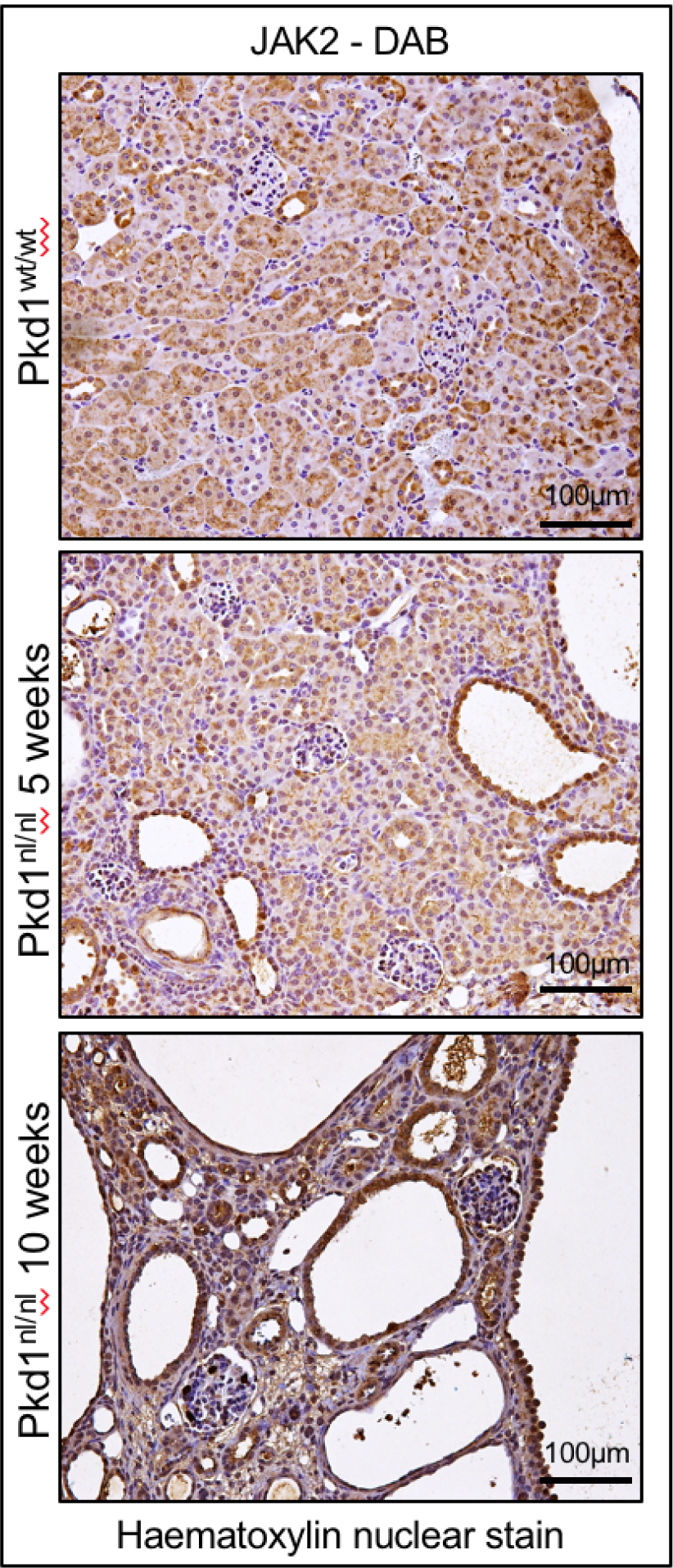
JAK2 is highly expressed by renal cystic epithelial cells *in vivo*. Kidneys sections of an orthologous ADPKD murine model, Pkd1^nl/nl^, at 5 and 10 weeks of age (5 mice in each group) were used to stain for JAK2. Immunohistochemistry of JAK2 was performed and representative images at 20x magnification are shown. The nuclei are counterstained with haematoxylin. Scale bars are 100μm.

### STAT3 tyrosine phosphorylation is potently suppressed by curcumin

Since JAK2 is highly expressed by cystic epithelial cells *in vivo* (Fig 1) we wondered whether its inhibition may be of therapeutic benefit and we set out to explore this biochemically in an *in vitro* setting. Studies in non-renal systems have suggested that curcumin, which is a natural phytochemical with strong anti-inflammatory activity and no appreciable side-effects, might be a JAK-STAT inhibitor ^18-20^. We therefore studied the ability of curcumin to inhibit JAK/STAT in three independent ADPKD-derived epithelial cells, three lines were used to avoid cell line specific artefacts. All ADPKD cells we have tested so far do not show signs of constitutive activation of JAK/STAT, therefore to activate the pathway we used a constant concentration of oncostatin M (OSM) ^21^. 50•• of curcumin for 6 hours potently inhibited the OSM-induced STAT3 activity, as evidenced by a marked reduction in tyrosine phosphorylation levels in all three lines tested (Fig 2A), without affecting total STAT3 protein levels. We next performed a time-response curve, which showed that one hour of curcumin treatment is sufficient to cause approximately 50% reduction in STAT3 phosphorylation while 3 and 6 hours cause nearly 100% reduction, while total STAT3 protein levels remained unaltered (Fig 2B). To study the concentration of curcumin required to cause 50% reduction in phosphorylation of STAT3 we stimulated cells for three hours with 100, 50 or 25•M of curcumin. We found that 100 and 50•M cause 100% reduction while 25•M cause about 50% reduction. These time- and dose-dependent data show that curcumin is a potent inhibitor of STAT3 activity.

**Figure 2:**
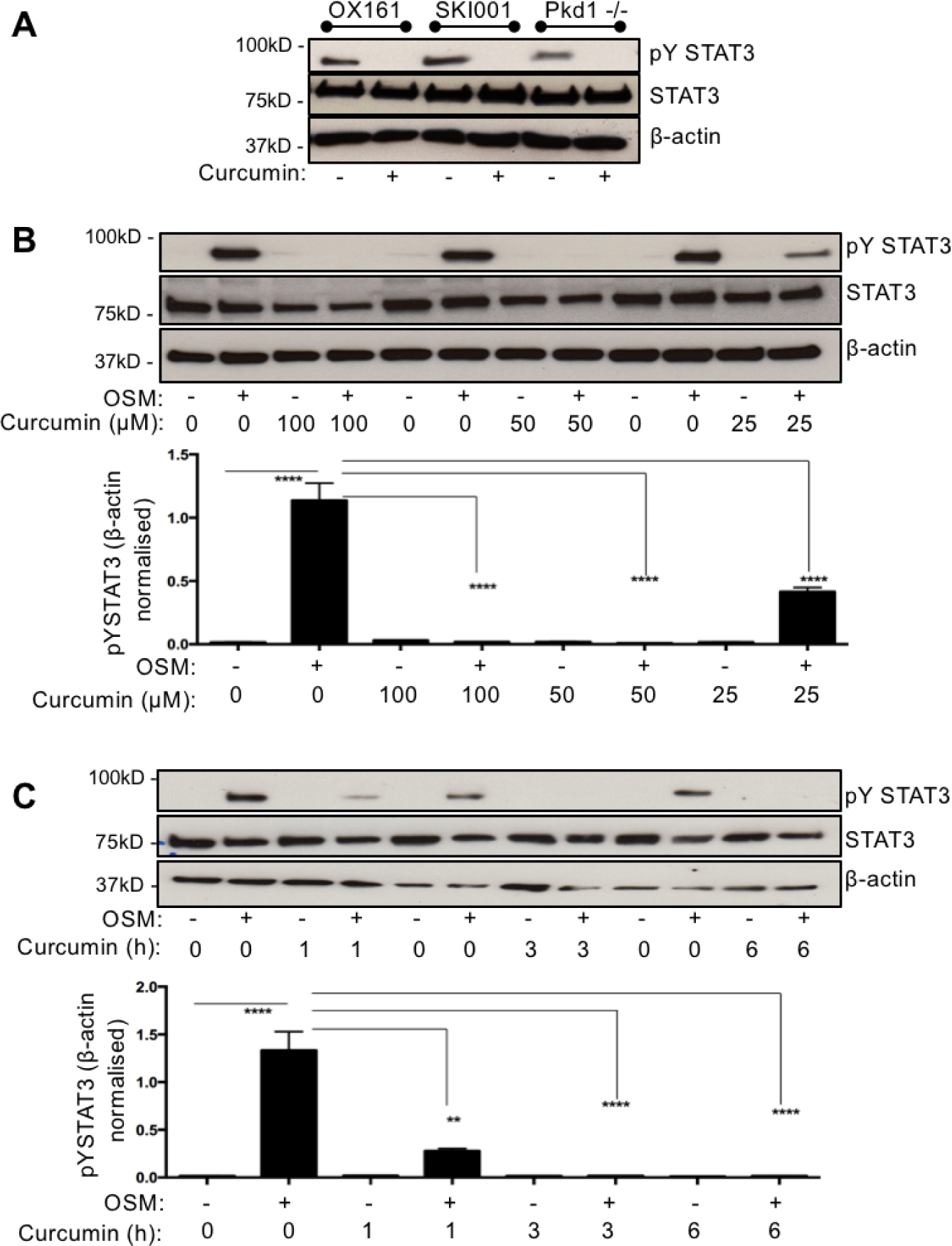
Curcumin is a potent STAT3 inhibitor. A. Oncostatin M (OSM, 10ng/ml) was used to stimulate phosphorylation of STAT3. Western blotting was performed (in human ADPKD-derived epithelial cells OX161, SKI-001 and mouse F1 Pkd1^-/-^) anti-STAT3 phospho-tyrosine 705 antibody (pY STAT3), total STAT3 (STAT3) and β-actin which served as an internal loading control were used. **B.** SKI001 cells treated with either 100, 50 or 25μM of curcumin were subjected to blotting for STAT3, pYSTAT3 and β-actin control. Densitometric quantification of three independent blots was carried out. One-way Anova with Bonferroni corrections was performed, and values lower that 0.05 were considered statistically significant. C. SKI001 cells were treated with 100μM of curcumin for 1 or 3 or 6 hours and stimulated either with water vehicle control or 10ng/ml of OSM. pYSTAT3, STAT3 and β-actin were investigated by blotting. Quantification of three independent blots was carried out. One-way Anova with Bonferroni corrections was performed, and values lower that 0.05 were considered statistically significant. Symbol meaning: * P≤0.05, ** P≤0.01, *** P≤ 0.001, **** P≤0.0001.

### Curcumin inhibits STAT3 activity via interfering with JAK2

Curcumin was shown by us and others to be an inhibitor of STAT3 phosphorylation, without altering STAT3 levels. An important question for any drug is to understand its mechanism of action. To identify how curcumin inhibits activity of STAT3 phosphorylation we studied JAK2, which is responsible for phosphorylating STAT3 and is highly expressed by cyst-lining cells in vivo (Fig 1). We studied JAK2 protein levels by immunoblotting and found that curcumin at 100 and 50•M caused a nearly 100% reduction in JAK2 protein levels, while 25•• of curcumin led to approximately 50% reduction (Fig 3). Interestingly curcumin reduced JAK2 levels in both OSM-treated and vehicle control-treated cells, therefore indicating that curcumin affects JAK2 in an OSM-independent manner in ADPKD cells. Curcumin’s effect on JAK2 activity was consistent with the inhibition of STAT3 phosphorylation observed above (Fig 2). Thus, these data suggest that curcumin inhibits JAK2 and thus reduces STAT3 activity.

**Figure 3:**
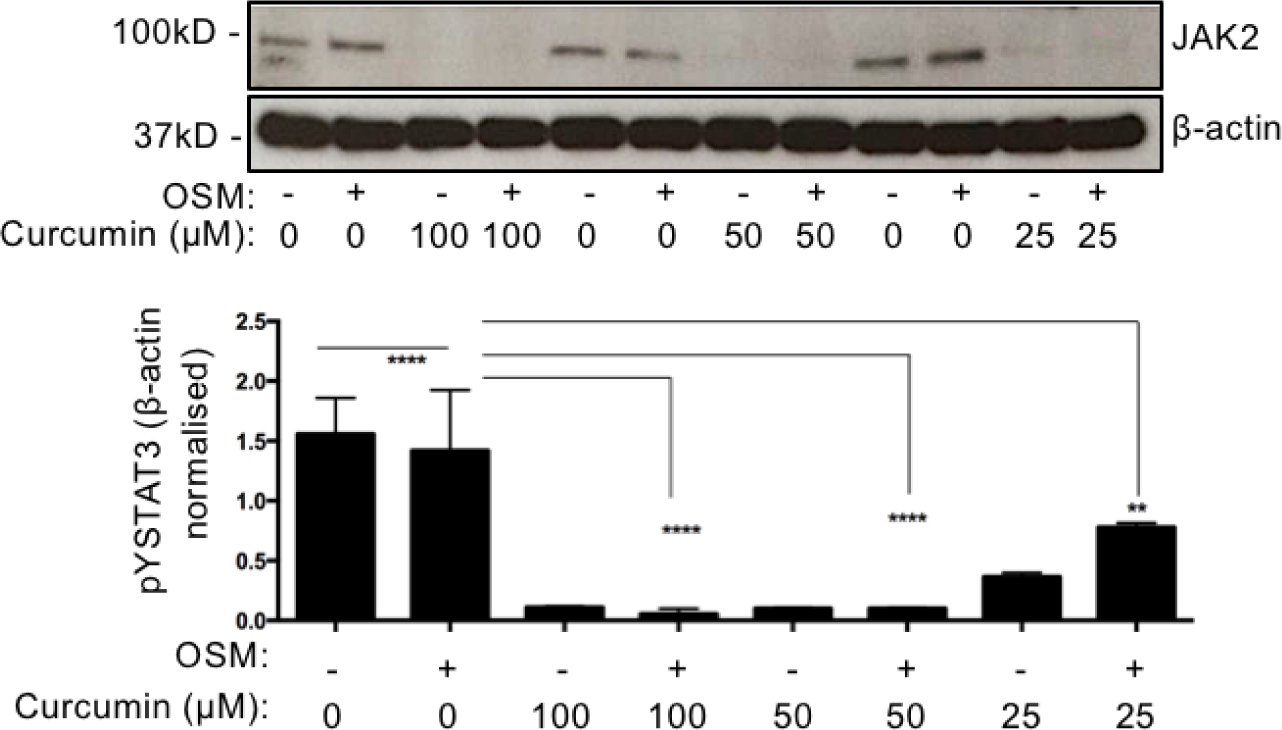
Curcumin inhibits JAK2. Western blotting of SKI001 lysates exposed to 100 or 50 or 25μM of curcumin was carried out, anti-JAK2 and β-actin were blotted. Quantification of three independent blots was carried out. One-way Anova with Bonferroni corrections was carried out and values lower that 0.05 were considered statistically significant. Symbol meaning: * P≤0.05, ** P≤0.01, *** P≤ 0.001, **** P≤0.0001.

### Curcumin does not change JAK2 levels but makes JAK2 insoluble

We wished to understand how curcumin causes a statistically-significant reduction in JAK2 levels. Firstly, we explored the possibility that curcumin may cause JAK2 to become proteosomally-processed. To study this, we started by treating cells with MG132, which is a proteasome inhibitor. We found as expected that MG132 caused an induction in ubiquitinated proteins (as it blocks their degradation), thus confirming that the proteasome was blocked sufficiently (Fig. 4A). However, curcumin reduced the amount of immunoblotted JAK2 levels in both OSM-treated and control conditions even when the proteasome was inhibited with MG132 (Fig. 4A compare lane 3 with 7 and 8). These data therefore indicate that the reduction in JAK2 is not mediated via proteosomal processing. Given that the proteasome is not responsible for the decreased levels of JAK2 we decided to perform immunocytochemistry to visualise JAK2 and see how curcumin treatment affects its cellular localisation. Interestingly, we found that JAK2 is localised throughout the cell body with some localisation in the plasma membrane in DMSO-vehicle treated cells, however once curcumin is added JAK2 becomes punctate (Fig. 4B). Previous studies have suggested that curcumin can cause protein precipitation into aggresomes ^22,23^, we therefore suggest that these punctate structures are JAK2 aggregates. This explains indicates that JAK2 become insoluble and therefore explains why we were unable to detect JAK2 in the detergent-soluble fraction by immunoblotting. Taken together, our data suggests that JAK2 is inactivated without being proteosomally-processed by curcumin.

**Figure 4:**
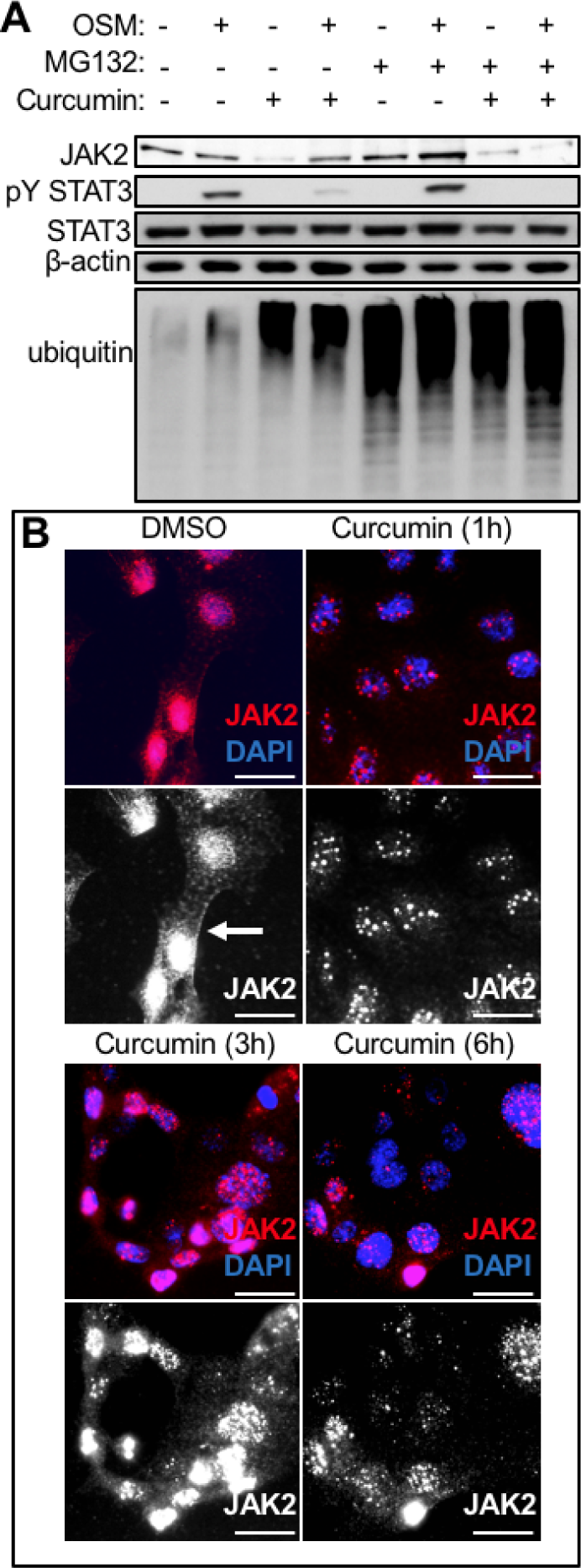
Curcumin controls JAK2 localisation. **A.** SKI001 cells were treated with OSM (10ng/ml), or MG132 (50μM) or curcumin (50μM) for 3 hours and lysates were subjected to immunoblotting. Antibodies against JAK2 pYSTAT3, STAT3, ubiquitin (indicator of proteasome inhibition control) and β-actin (internal loading control) were studied. **B.** SKI001 cells were either treated with DMSO vehicle control for 3 hours (3h) or curcumin for 1h or 3h or 6h, they were then snap frozen on ice-cold methanol, stained with anti-JAK2 rabbit antibody, followed by an anti-rabbit AF594 and imaged in an inverted fluorescent microscope. Red is anti-JAK2 (also in greyscale in lower panel), blue is nuclear counterstain (DAPI). Scale bars are 25μm.

### Curcumin-mediated JAK2 inhibition reduces cystic growth *in vitro*

To study the ability of the JAK2/STAT3 pathway to drive cystogenesis we performed organotypic cyst assays *in vitro*. We used three independent cell lines to ensure robustness of results and avoid cell-type specific artefacts. In these assays microscopic cysts form and grow in diameter over time, as previously shown ^5^. We found that curcumin-induced JAK2 inhibition led to a statistically-significant and consistent reduction in the size of cysts in all three independent cell lines tested and, in a time and dose-dependent manner (Fig 5). Specifically, curcumin caused a statistically-significant blockade in growth of cysts already at day 2 at the highest concentration, which persisted to day 6 and 9, but with 50 and 25•• of curcumin also significantly reducing the cyst size by day 9 in OX161 cells, which are human-derived ADPKD cells (Fig 5A). A similar pattern was seen in SKI-001, an additional independent human ADPKD-derived line, which showed reduced cyst size at day 6 and 9 with 50 and 25•• of curcumin (Fig. 5B). Finally, we also studied a mouse line which has a Pkd1 deletion and is therefore considered as a mouse ADPKD line, as expected curcumin blocked the growth of cysts at day 2 with 100•M of curcumin, while day 6 and day 9 50 and 100•• were able to cause a block in the growth of cysts (Fig. 5C). Taken together, these data provide first evidence that inhibition of JAK2/STAT3 can modulate cystogenesis *in vitro*.

**Figure 5:**
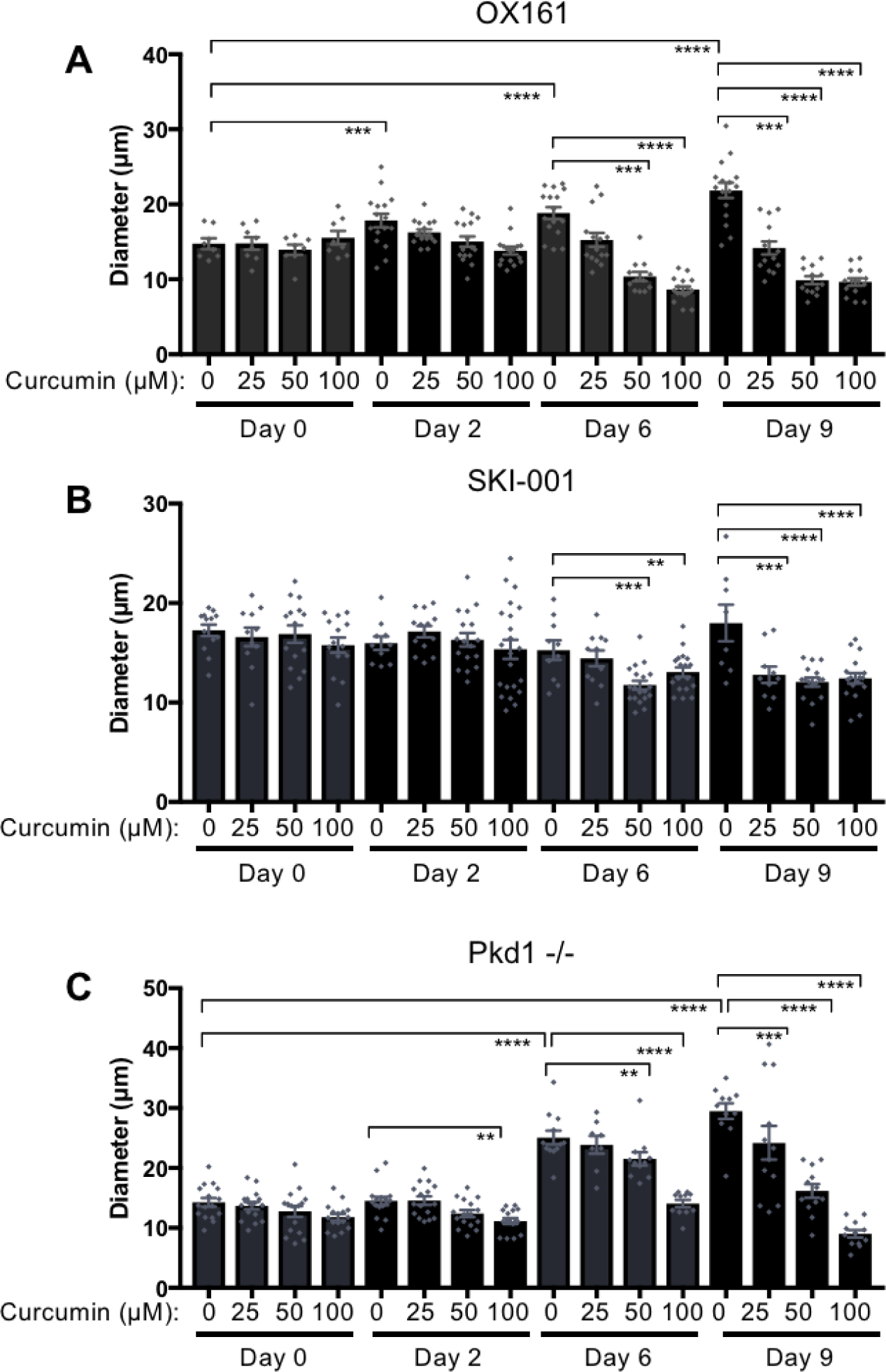
Curcumin slows down cyst growth *in vitro*. ADPKD-derived human epithelial cells were grown in BD-Matrigel to allow the formation of cysts to grow over time. All cysts formed were photographed at day 0, 2, 6 and 9; using an Olympus inverted microscope. Curcumin at 100, 50, 25 and 10μM or DMSO vehicle-control was added and cyst growth measured in **A** human OX161 **B** human SKI-001 and **C** mouse derived Pkd1^-/-^ cell lines. One-way Anova with Bonferroni corrections was carried out and values lower that 0.05 were considered statistically significant. Symbol meaning: * P≤0.05, ** P≤0.01, *** P≤ 0.001, **** P≤0.0001.

## Discussion

This is the first study to identify strong JAK2 expression in cysts of the ADPKD mouse kidney *in vivo*. Moreover, we are the first to demonstrate that suppression of JAK2 leads to blockade in cystogenesis *in vitro*. We therefore propose that JAK2 is a key signalling tyrosine kinase in the polycystic kidney and this study provides the foundation for the use of JAK2 inhibitors as therapeutics in ADPKD. Indeed, a number of groups, including our own, have shown that STAT3, STAT6 and STAT5 are miregulated in ADPKD ^5,8,9,24-29^. Interestingly, JAK2 can indeed activate STAT3 ^30^ STAT5 ^31^ and STAT6 ^32^. Moreover, JAK2 plays a central role in maintaining cell proliferation ^33^ and its inhibition suppresses proliferation ^34^, which is a key misregulated function in ADPKD. Our work therefore suggests that JAK2 upregulation is responsible for the abnormal activation of STATs in the polycystic kidney. On the contrary, it has been suggested that STAT3 activation can be JAK2-indepenenent and instead Src-mediated in ADPKD ^29^; interestingly curcumin has been shown to inhibit Src-mediated signalling ^35,36^. Therefore, future work using selective kinase inhibitors is needed to decipher the pathways that participate in disease development.

Over the years, phytochemicals have been used to inhibit, prevent or reverse disease processes. Curcumin specifically was found to have strong anti-cystic effects in murine models of ADPKD ^16^. Whether curcumin acts primarily on renal epithelial or vascular cells is however unclear, making the interpretation of the *in vivo* results ^16^ difficult. Moreover, extra complications arise due to the fact that curcumin is currently under trial (phase 4) for its potential ability to modify the vascular phenotype in children and young individuals with ADPKD. Our work identifies that curcumin directly affects renal tubular epithelial cells, without requiring interactions with the vasculature, thus providing some answers. However, expression studies show that JAK2 is also expressed by vascular cells, thus not precluding a role of curcumin in the vasculature in ADPKD.

The use of curcumin as a therapy has both advantages and disadvantages. One advantage is the possibility of long term use, due to lack of appreciable side effects ^37^. However, curcumin’s reduced bioavailability ^38,39^, makes it less attractive when compared with small molecule inhibitors. Indeed, our study identifies JAK2 as a therapeutic target in ADPKD. Given that JAK2 inhibitors exist, an interesting development would be to compare and contrast newest selective JAK inhibitors with older generation pan-JAK inhibitors and phytochemicals such as curcumin.

Overactivation of JAK2 can lead to malignancy ^40^. In our study we find that expression of JAK2 is wide spread in cortical tubules, however no dysplastic phenotype in seen ADPKD, with cysts arising from nephrons that otherwise look normal. This suggests that activation of JAK2 in ADPKD leads to a change in the maintenance of epithelial differentiation and turnover. This is supported by our previous work which showed that increased JAK-STAT activity led to enhanced proliferation in renal tubular epithelial cells ^5^. Whether inhibition of JAK activity leads to renal protection in ADPKD via reducing tubular epithelial cell growth and or cell death will need to be investigated in future studies.

Intriguingly, curcumin causes JAK2 to move into an insoluble fraction consistent with generation of multi-molecular structures or aggresomes. Therefore, we propose a model whereby curcumin takes JAK2 temporarily out of action by causing it to aggregate, without causing JAK2 to degrade which is an irreversible consequence. Temporarily inhibiting JAK2 may explain why curcumin is not cytotoxic, allowing JAK2 to become reactivated when needed. Likewise, others have reported that curcumin-treated cancer cells respond by inhibiting STAT3, a curcumin-induced effect which is fully reversible ^41^. How curcumin causes abnormally activated JAK2 to aggregate is currently unknown. Curcumin has been shown to be neuroprotective, by preserving gross intramolecular conformation of beta amyloids, while disrupting the intermolecular arrangements, a phenomenon which is very similar to the effects of Zn^2+^ ions, as reported previously ^23^. Likewise, curcumin can alter the structure of lipoproteins by interacting with the soluble intermediates, thus altering their aggregation patterns ^22^. Similarly, A20, which is an anti-inflammatory zinc-finger protein is shown to cause aggregation of NF•B (an inflammatory transcription factor) and mediate vascular protection ^42^. Based on previous studies we therefore hypothesize that curcumin disturbs JAK2 intramolecular interactions using a similar mechanism.

Given that the understanding of the mechanisms underlying cystic growth can provide crucial insights for the development of therapeutic agents, the identification of molecules that play important roles in proliferation and cystogenesis is therefore imperative. Given the novelty of using JAK inhibitors to treat ADPKD more studies need to be carried out to further understand the mechanisms employed by JAK-STAT to regulate cystic growth. Nonetheless, our study suggests that JAK2 is a key signalling molecule in ADPKD and its inhibition may provide the foundation for its development as a novel therapeutic in polycystic kidney disease.

## METHODS

### Cell lines

Conditionally-immortalized ADPKD-derived lines (OX161 and SKI-001) ^43^ are tubular epithelial cells isolated from human kidneys and immortalized by transduction at an early passage (P1-4) with a retroviral vector containing a temperature-sensitive large T antigen and the catalytic subunit of human telomerase ^44^. The F1-Pkd1 WT renal epithelial cells were previously isolated from kidney papillae of the Pkd1fl/fl mouse (B6.129S4-Pkd1tm2Ggg/J; the Jackson Laboratory) and were immortalized with the lentiviral vector VVPW/mTert expressing the mTert; to delete Pkd1 and produce F1/Pkd1-/- cells they were subsequently transfected with VIRHD/HY/Silnt•1/2363 lentivectors followed by hydromycin selection ^45^.

### Cyst assays

Cyst assays were performed as previously described ^5^. In brief, for ADPKD cystic cell lines, 2 human lines namely OX161, SKI-001, and one mouse line F1-Pkd1^-/-^ were used. OX161 and SKI-001 were kept at 33°C F1-Pkd1-/- were at 37°C. Healthily growing cells were trypsinised and re-suspended in DMEM media containing 10% FBS (no antibiotics) and counted using a haemocytometer. Subsequently 2×10^4^ cells per well were mixed with basement Matrigel (354230, BD biosciences). Prior to use the Matrigel was left to thaw on ice and once cells were added it was immediately loaded onto the 96 well plate, allowing it to polymerise. 100•l of DMEM media supplemented with 10% FBS was added to each well and cells returned to cell incubator for 24 hours to allow them to recover. After 24 hours, the media was aspirated and replaced with fresh media (DMEM + 10% FBS) supplemented with either vehicle (DMSO) or curcumin (Sigma). The cells were photographed before every media change and every two days, the media was replaced with fresh media containing either curcumin or DMSO.

### Immunohistochemistry/immunofluorescence

Cells grown on coverslips were fixed with ice-cold methanol prior to blocking in 2% milk/TBST for 30 minutes and incubated with primary antibodies overnight at 4°C. Antibodies used was anti-rabbit JAK2 (3230S, cell Signalling, D2E12). Cells were washed and incubated with secondary which was an anti-rabbit AF594 (A-11037, Invitrogen (1:250) antibody. Microscopy was carried out using a confocal microscope. Immunohistochemistry using a JAK2 antibody (3230S, Cell Signalling) was performed as previously described ^5^.

### Western blotting and curcumin treatment

Cells were analysed by Western blotting with previously reported protocols ^46-49^. In brief, cells were lysed using ice-cold Lysis Buffer (50mM Tris (pH 7.4) 250mM NaCl, 0.3% Triton X-100, 1mM EDTA) supplemented with protease inhibitor cocktail (Roche), freeze-thawed and sonicated. Whole cell lysates were boiled in 2xLaemmeli sample buffer for 5 minutes. Samples were resolved by SDS-PAGE and transferred using the Mini-PROTEAN system (Bio-Rad). Primary antibodies were anti-•-actin (ab8226, Abcam), anti-phosphorylated STAT3 (p-STAT3) (9145, Cell Signalling), anti-STAT3 (12640, Cell Signalling). Curcumin (08511-10MG, Sigma) was resuspended in DMSO and used at stated doses, DMSO served as vehicle control.

### Animals

*Pkd1*^nl/nl^, harboring an intronic neomycin-selectable marker, or wild type littermate controls were used ^5,50^. *Pkd1*^nl/nl^ or control mice were sacrificed at 5 or 10 weeks of age and kidneys collected, formalin-fixed and paraffin embedded. 5-micron section were used for histological examination. All mouse experiments were done under the authority of a U.K. Home Office license.

### Statistical analysis

Data were analysed using Prism GraphPad and Non-parametric, two-tailed, Mann-Whitney T-tests or one-way ANOVA were performed. Results with a P value of 0.05 or lower were considered statistically significant.

## Acknowledgements

We would like to thank Prof AC Ong and his group for providing advice and cell lines; Ms Fiona Wright for tissue processing; Ms Monica Neilan for genotyping, Prof DJ Peters and her group for the Pkd1^nl/nl^ animal model^17^ and Luca Gusella for the collecting duct Pkd1^-/-^ F1 cells. This work was supported by a Kidney Research UK post-doctoral fellowship and a Women Academic Returner’s Programme (WARP) from the University of Sheffield awarded to Maria Fragiadaki.

